# Cell-penetrating peptide conjugates of indole-3-acetic acid-based DNA primase/Gyrase inhibitors as potent antitubercular agents against planktonic and biofilm culture of *Mycobacterium smegmatis*

**DOI:** 10.1101/2020.12.01.406405

**Authors:** Rikeshwer Prasad Dewangan, Meenakshi Singh, Stefan Ilic, Benjamin Tam, Barak Akabayov

## Abstract

*Mycobacterium tuberculosis* (*Mtb*) is a pathogenic bacterium that caused 1.5 million fatalities globally in 2018. New strains of *Mtb* resistant to all known classes of antibiotics pose a global healthcare problem. In this work we have conjugated novel indole-3-acetic acid-based DNA primase/gyrase inhibitor with cell-penetrating peptide via cleavable and non-cleavable bonds. For non-cleavable linkage, inhibitor was conjugated with peptide via an amide bond to the N-terminus, whereas a cleavable linkage was obtained by conjugating the inhibitor through a disulfide bond. We performed the conjugation of the inhibitor either directly on a solid surface, or by using solution-phase chemistry. *M. smegmatis* (non-pathogenic model of *Mtb*) was used to determine the minimal inhibitory concentration (MIC) of the synthetic conjugates. Conjugates were found more active as compared to free inhibitor molecules. Strikingly, the conjugate also impair the development of biofilm, showing a therapeutic potential against infections caused by both planktonic and sessile forms of mycobacterium species.

## 1. Introduction

The global development of antibiotic-resistant pathogens is currently one of the biggest challenges in healthcare (Laxminarayan et al., 2013). One specific example is the treatment of infections caused by *Mycobacterium tuberculosis* (*Mtb*) that became a serious threat due to the development of multidrug-resistant strains (e.g., extended and total drug-resistant pathogens) in both community and nosocomial environments (Seung et al., 2015). Even the last resorts of conventional anti-tubercular therapies are unable to treat these infections efficiently in patients. Also, the current development of new agents with a novel mode of action, is not sufficient to fight these superbugs (Fair & Tor, 2014).

*Mtb* has several survival mechanisms that render its treatment a challenge for contemporary medicine. Namely, this pathogen can survive and persist in macrophages by blocking the fusion of phagosomes with lysosomes (Flynn & Chan, 2001). Another extraordinary defence strategy of this organism is its capability to form biofilm under environmental stresses, including the presence of antibiotics (Ojha et al., 2008). It is, therefore, not surprising those 1.4 million deaths due to *Mtb* infections have been reported in 2015. It is also important to note that out of a total of 10.4 million reported cases, only ~10% of patients developed active tuberculosis (TB), whereas the remaining 90% exhibited latent infection. However, even this latent infection may convert into an active form at a certain time-point, especially if the immune system gets weakened (Liu et al., 2018; WHO & Regional Office for South-East India, 2017).

Conventional anti-tubercular therapy is based on a few classes of anti-tubercular drugs that usually target metabolic pathways of the bacterium (Rastogi & David, 1993). Therefore, the bacteria may mitigate the effect of these drugs by incorporating DNA mutations into genes that encode related protein targets. Another problem is that the upcoming new drugs are either analogues of conventional medicines or novel chemicals with the same targets. Hence, explorations of novel targets offering different treatment approaches are needed (Mantravadi et al., 2019). The discovery and development of bedaquiline (Sirturo) is a good example of a clinical therapeutic, with a novel target, which enables successful treatment of MDR-TB infections (Andries et al., 2005).

Replication of the chromosome is a central event in the cell cycle of every bacterium; it is performed by the replisome, a multi-enzyme complex that synthesizes DNA continuously at the leading strand and discontinuously at the lagging strand (Marians, 2008; Yao & O’Donnell, 2010). The bacterial replisome fulfils most of the requirements of a potential novel drug target: DNA replication proteins are essential for viability, and many of the DNA replication proteins present a different setting in bacteria than in humans, thereby providing high selectivity (Ilic et al., 2018). Therefore, inhibiting any of the essential enzymes associated with DNA synthesis should affect bacterial growth in planktonic and sessile forms of bacteria (Rose et al., 2015; Trivedi et al., 2016).

Using a newly developed NMR-fragment based virtual screening (FBVS), we previously discovered inhibitors that target DNA primase of bacteriophage T7, which serves as a model for bacterial DnaG-type primases (Ilic et al., 2016). Not surprisingly, some of these inhibitors also target DNA replication in *Mtb* by inhibiting simultaneously two enzymes, namely DnaG primase and DNA gyrase, since these two enzymes share a conserved structural domain near the inhibitors’ binding site. SAR studies were performed to improve the physicochemical properties of the molecules, as well as inhibitory activity against DnaG primase and DNA gyrase, resulting in novel indole acetic acid-based selective inhibitors (IN) (Figure 1). These exhibited bacteriostatic activity against *Mycobacterium smegmatis* (*Msmg*) (M. Singh et al., 2020). However, the potency of these molecules, as examined using *in-vitro* enzymatic assays was not high, and they were moderately active against *Msmg*. Furthermore, pathogenic *Mtb* presents an intrinsically impermeable thick cell wall (Hett & Rubin, 2008), which serves as a robust barrier to most antibiotics. Therefore, a major limitation for conventional antitubercular drugs against intracellular infections of *Mtb* is the ability to cross several biological membranes to reach designated biological targets within pathogens (Rohde et al., 2007).

**Figure: 1.**
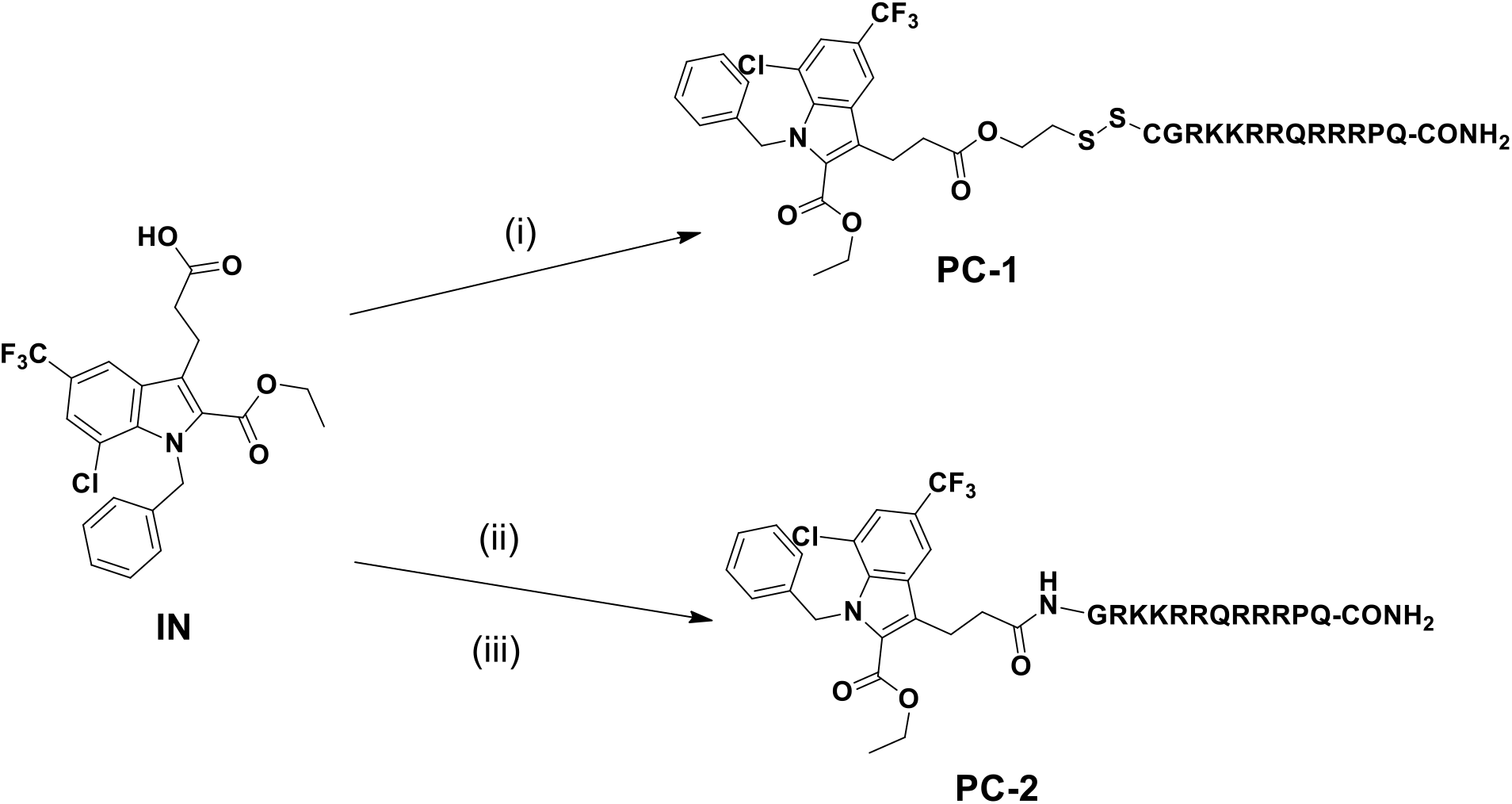
Structures and synthesis of peptide conjugates of DNA primase/gyrase inhibitor. Reagents and Conditions: i). SH-CysTAT-CONH2 MOPS Buffer, pH-7.4, 24h, at room temperature, ii) H2N-TAT-Resin, HBTU, DIPEA in DMF, iii) TFA: Thioanisole: EDT: Pheno1: water (82.5:5:2.5:5:5), 3h.

Intracellular delivery using membrane-permeable peptide vectors is a recently developed methodology that has been employed successfully to transport various bioactive molecules into mammalian and mycobacterium cells to modify cellular functions (Deshayes et al., 2005; Sparr et al., 2013). Arginine-rich peptides such as HIV-TAT were shown to be effective in translocation across the membrane (Mishra et al., 2011). Therefore, the development of novel strategies to improve the delivery and the bioavailability of existing or novel drugs is an opportunity to significantly increase efficacy and/or reduce systemic toxicity through dose reduction (Jean et al., 2016).

It has been reported that TAT and their analogues show inherent antimicrobial activities or enhancement of antimicrobial activity of other molecules. For example, TAT peptide dimers (Long Zhu & Shin, 2009), TAT peptide-porphyrin conjugates (Bourré et al., 2010) and D-TAT (Jung et al., 2008) (diastereomeric analogue) were reported to have antibacterial activity against gram-positive and gram-negative bacterial strains. Self-assembled cholesterol-conjugated Tat peptides (Wang et al., 2010) were shown to form nanoparticles and exhibit antifungal activity against human pathogenic fungi. Bhosle et al. reported the activity of TAT peptide and their peptidomimetic analogues against a broad range of bacterial strains and *Mtb* in dormant and active forms. This opens the scope of further development in design of TAT or their conjugates (Bhosle et al., 2018).

The broad application of cell penetrating peptides as antimicrobial agents motivated us to investigate the effect of conjugation of TAT peptides with our novel DNA primase/gyrase inhibitor (IN) against the tuberculosis model of *Msmg*. TAT peptides were conjugated with IN via amide and disulfide linkages. Then we evaluated the *in vitro* DNA primase/gyrase inhibition activity of these peptide conjugates as well as their efficiency against the growth of *Msmg* mc^2^155. In addition, we evaluated the inhibition of biofilm formation induced by these conjugates.

## 2. Results

### 2.1 Synthesis and characterization of peptides conjugates

The design, synthesis, and optimization of DNA/Gyrase inhibitor (IN) were reported in our previous work (M. Singh et al., 2020). Within the series reported in our previous work we have selected most active analogue (IN) for conjugations. The characterization of the compound IN is provided in the supporting information (ESI). Conjugation with a cell-penetrating NH2-TAT-CONH2 peptide (NH2-GRKKRRQRRRPQ-CONH_2_) was performed via disulfide (PC-1) and amide bonds (PC-2). The TAT peptide was synthesized by 9-fluorenyl methoxycarbonyl (Fmoc) based solid-phase peptide synthesis (SPPS), and the sequence was characterized by analytical RP-HPLC and ESI-MS (Table 1 and Figure S1-S8).

**Table 1:**
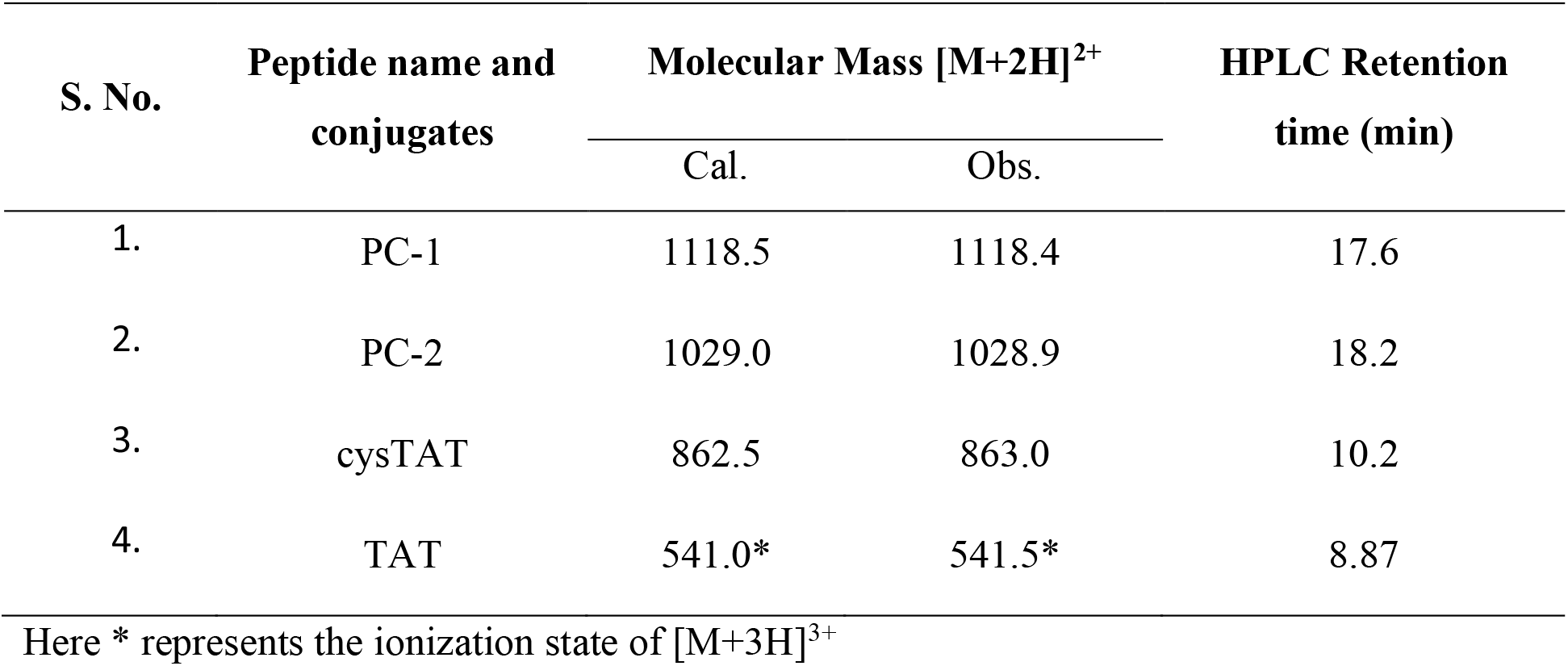
Sequence, HPLC and MS characterization of synthesized peptide conjugates

### 2.2 Inhibition of *M. smegmatis* growth

We have conjugated a novel inhibitor, IN with cell-penetrating peptides (Figure 4c): HIV TAT (GRKKRRQRRRPQ-CONH2) and CysTAT and evaluated their ability to inhibit growth of *Msmg* mc^2^155. *Msmg* mc^2^155, a non-pathogenic, fast-replicating mycobacterium, is widely used as a model system to study *Mtb* (McGuire et al., 2012; Mohan et al., 2015). *Msmg* mc^2^155 and *Mtb* share significant similarities in their genomes, including genes associated with mycolic-acid rich cell walls (A. K. Singh & Reyrat, 2009).

**Figure: 3.**
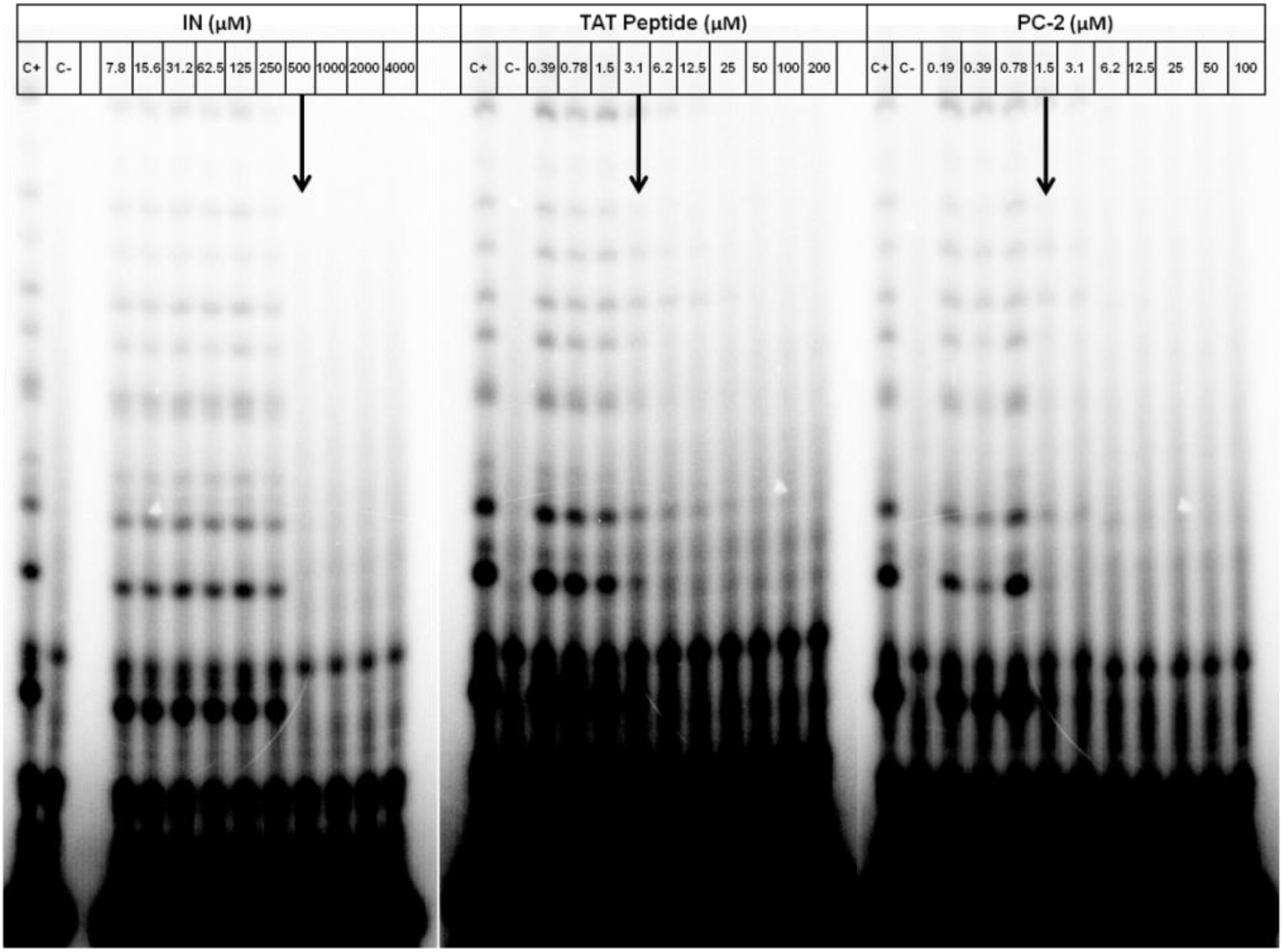
Effect of Inhibitor molecule (IN), PC-2 and TAT peptide on RNA primers synthesis catalyzed by DNA primase from *Mtb*. The reaction contained the oligonucleotide 5’-CCGACCCGTCCGTAATACAGAGGTAATTGTCACGGT-3’, α-32P-ATP, CTP, GTP, UTP in a standard reaction mixture, and different concentrations in double dilutions of each small molecule. The reaction was initiated by adding *Mtb* DnaG primase to a final concentration of 500 nM. After incubation, the radioactive products were analyzed by electrophoresis through a 25% polyacrylamide gel containing 7 M urea and visualized using autoradiography. Indicated arrow shows the concentrations of the inhibitor molecules, peptide and conjugates at which inhibition of the enzymes was occurred.

**Figure: 4.**
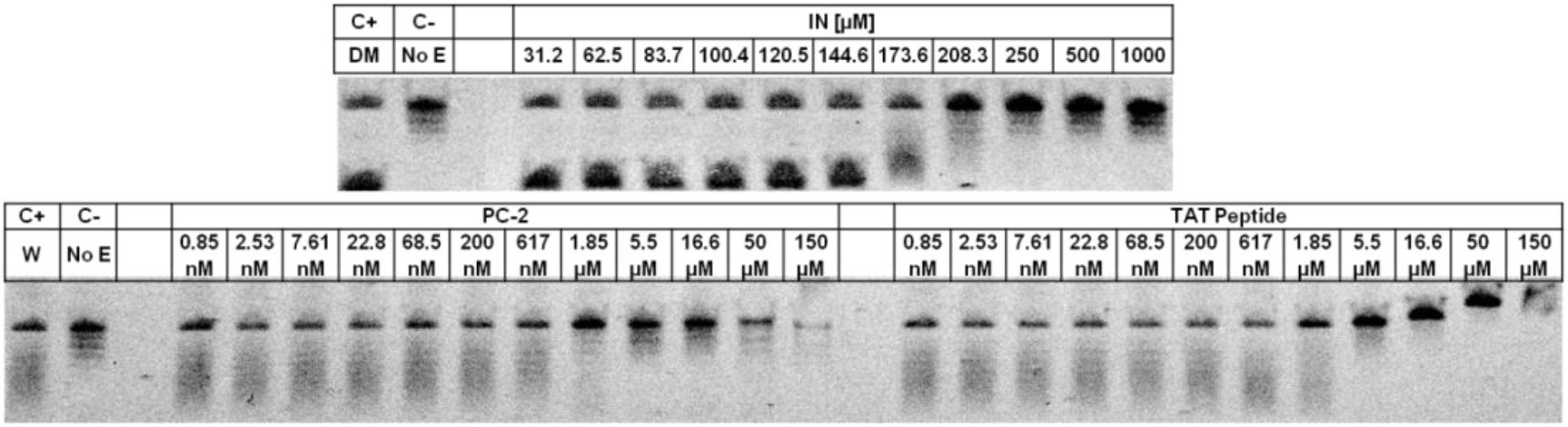
Effect of inhibitor molecule (IN), PC-2 and TAT peptide on inhibition of DNA supercoiling by small molecule inhibitors. The 20μL reaction containing DNA Gyr (reconstitution of 36 nM GyrA and 18 nM GyrB), l3.24ng/μL pBR322 DNA and 1 mM ATP in the presence of inhibitors was incubated at 37 °C for 60 min. The reaction products were resolved in 1% agarose gel followed by staining with EtBr.

Interestingly, the conjugated molecules were found to be >50 fold active against the *Msmg* mc^2^155, compared to compound IN alone, with the MIC range of 1.9 – 3.9 μM (Table 2). Under identical conditions, we also evaluated the antibacterial activity of cell-penetrating peptides, and MIC was found to be 7.8 μM for both conjugated peptides (Table 2). Hence, we have concluded that the conjugation of our experimental inhibitor to these peptides facilitated the cellular uptake and potentiated the antibacterial activity.

**Table 2:** Minimum inhibitory concentrations (MICs) of the inhibitor, peptide conjugates and free peptides against *M. smegmatis* mc^2^155

| S. No. | Name of compounds | MIC<sub>90</sub> (μM) |
| --- | --- | --- |
|  |  | *M. Smegmatis* mc<sup>2</sup>155 |
| 1. | IN | 250 |
| 2. | PC-1 | 3.9 |
| 3. | PC-2 | 1.9 |
| 4. | TAT | 7.8 |
| 5. | Cys-TAT | 7.8 |
| 6. | Isoniazid | 250 |

### 2.3 *In vitro* enzyme inhibition assays

#### 2.3.1 Inhibition of DnaG activity

*Mtb* DNA primase (DnaG) has been reported as a potential drug target to design new anti-tubercular compounds to fight drug-resistant TB. We have designed indole-3-acetic acid based novel compounds which have shown inhibitory activity against DnaG (M. Singh et al., 2020). Next, we wanted to explore if the inhibitory activity would be improved upon conjugating these molecules to a peptide carrier. The protocol for *in vitro* DnaG-catalyzed RNA primer synthesis was reported previously (M. Singh et al., 2020). Briefly, this reaction involved incubation of *Mtb* DnaG with DNA template (5’-CCGACCCGTCCGTAATACAGAGGTAATTGTCACGGT-3’), along with rNTPs, and [α-^32^P]-ATP. Due to enzymatic activity of DnaG, radiolabelled RNA primers were synthesized. The inhibitor (IN) and conjugates were added into the reaction mixture to investigate their ability to impair oligoribonucleotide synthesis by *Mtb* DnaG. The RNA products were separated by denaturing gel electrophoresis and visualized on an autoradiogram (Figure 3). The decrease in the intensity of radio-labelled RNA primers, upon increasing concentration of the components and the conjugate (IN, TAT peptide, and PC-2, respectively), indicates inhibition of the *Mtb* DnaG. Here, we used the most active conjugate PC-2 and its corresponding peptide for enzyme inhibition assay. Peptide conjugates significantly increase inhibition of *Mtb* DnaG compared to unconjugated small molecules (Figure 3). While the compound IN inhibits the enzyme activity at concentrations of 500μM, peptide conjugate of this molecule inhibits at concentrations of 1.5 μM. The inhibitory concentration of these molecules is proportional to their calculated MIC against *Msmg* (Table 2).

#### 2.3.2 Inhibition of gyrase activity

To assess the activity of Gyr in the presence of experimental inhibitors DNA supercoiling assays were carried out (Figure 4). In the absence of inhibitor, the enzyme showed marked DNA relaxation activity in the presence of ATP, observed *in vitro* as negative supercoiling of pBR322 DNA. After the incubation, the reaction products were separated and analysed by gel-electrophoresis. The reduction of supercoiling activity resulted in DNA band shift or the appearance of multiple bands relative to the control sample (Figure 4). The compound IN was able to inhibit the supercoiling of DNA at concentrations of 250μM, whereas peptide conjugate (PC-2) of this molecule was able to inhibit at concentrations of 1.85 μM. The TAT peptide was also evaluated for inhibition of supercoiling activity of the gyrase enzyme and found inhibitory concentrations 5.5 μM, which was a 3-fold higher concentration than the value observed for PC-2. Interestingly, the inhibitory concentrations of these molecules correspond to their calculated MIC against *Msmg* (Table 2).

### 2.4 *In vitro* inhibition of biofilm formation by *Msmg*

Previous studies have shown that inhibition of DNA replication affects biofilm formation and stability of preformed biofilm. This link between DNA replication and biofilm formation is likely to be consequential (Gotoh et al., 2008). To evaluate the effect of small-molecule inhibitor (IN) and its peptide conjugates on *Msmg* biofilm formation, cells were treated with each compound and the amount of biofilm formed was quantified as reported previously (Anand et al., 2015). Initial inoculums of *Msmg* were incubated with MIC and sub-MIC of experimental inhibitors, including CPP conjugates, and isoniazid as a control. At MIC (250 μM) and sub-MIC (125 μM) IN inhibited biofilm formation up to 73.5% and 11.2%, respectively, as compared to DMSO control (Figure 5a). The PC-2 conjugate at MIC and sub MIC level was found to inhibit biofilm formation up to >99%. In identical conditions the peptides alone were not able to inhibit the biofilm formation in both concentrations. Surprisingly, the standard drug isoniazid reduced biofilm formation up to 66% at MIC and did not inhibit the biofilm formation significantly at sub-MIC level (Figure 5a). To further corroborate these results, the alamar blue cell viability assay was performed. In addition, we also counted the CFU from *Msmg* biofilm suspensions treated with our experimental compounds by plating each sample on 7H10 agar plate (data has not shown).

**Figure: 5.**
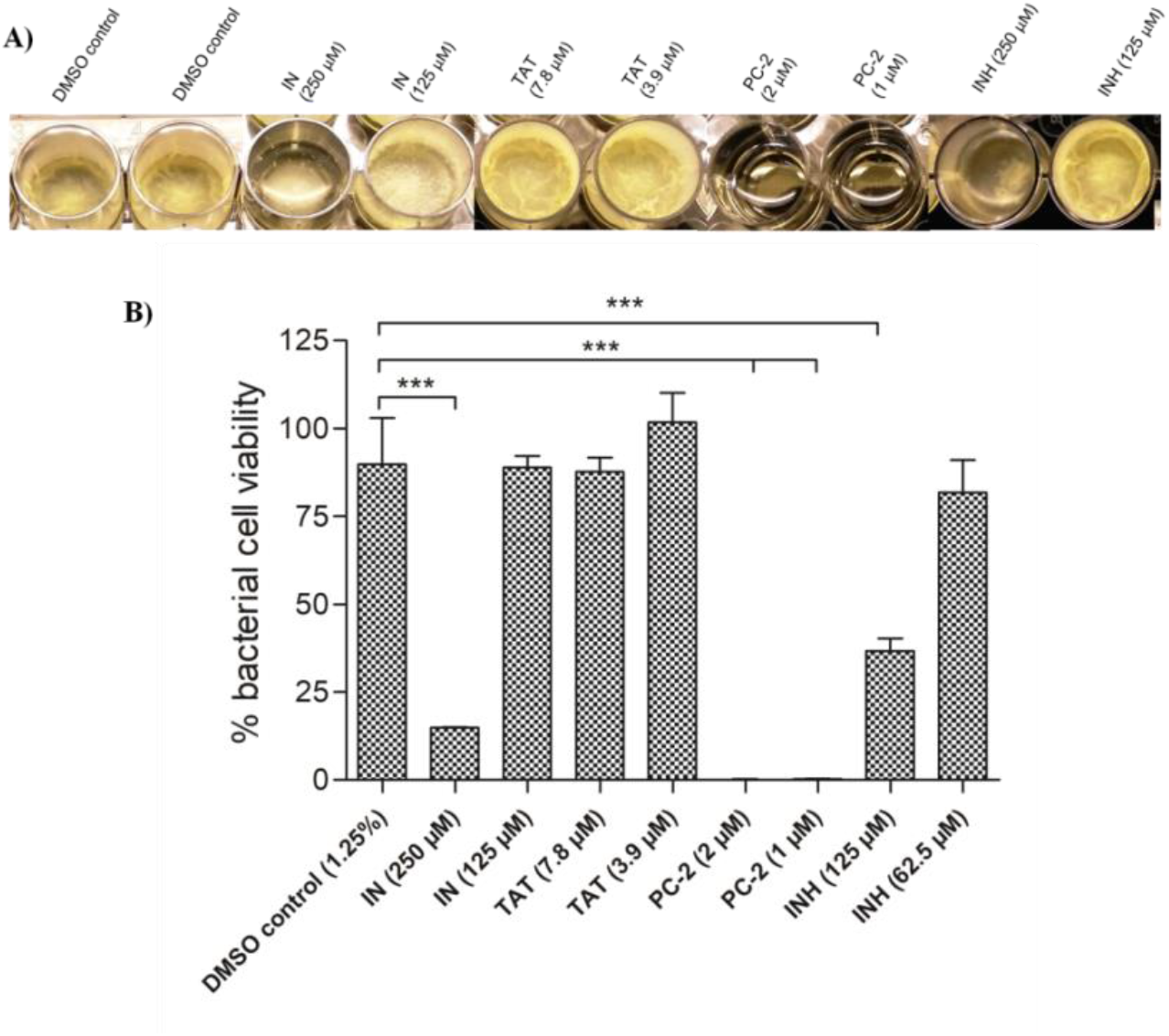
Inhibition of *Msmg* mc^2^155 biofilm formation. A) Optical images of *Msmg* biofilm with or without treatment with experimental inhibitors B) Quantification of bacterial cell viability in biofilm formation inhibition assay.

## 3. Discussion

Antibiotic resistance is a global concern, especially in bacteria that possess unique survival mechanisms, such as mycobacteria (Kamaruzzaman et al., 2017). For example, tubercle bacilli can remain in viable form for many years in the host, due to their specific cell wall composition as well as the ability to hide inside macrophages (Lerner et al., 2015). Therefore, the discovery of new molecules with novel modes of action is required to treat emerging drug-resistant tuberculosis infections. The goal of this study was to investigate the influence of chemical linkage between novel DNA primase/gyrase inhibitor and TAT peptide.

DNA primase is an essential component of DNA replication that synthesizes short RNA primers, used by the DNA polymerase to form “Okazaki fragments” on the lagging DNA strand (Marians, 2008). On the other hand, the gyrase is employed to release the twisting tension generated by DnaB helicase during DNA unwinding. Gyrase acts by creating a transient double-stranded DNA break and catalyzing the negative supercoiling of DNA at the expense of ATP hydrolysis (Keszthelyi et al., 2016). Both DnaG primase and Gyr are essential for viability, and have different structural and functional settings in human replisomes, thereby providing attractive candidates for antibiotic targeting (Ilic et al., 2018). Importantly, these enzymes in prokaryotes share common TOPRIM fold that constitute the side wall of the binding-site cleft of the small molecule inhibitors we found. Using NMR-fragment based screening and computational optimization we found novel indole-3-acetic acid derivatives that inhibit T7 primase but also DnaG and Gyrase of *Mtb* (Ilic et al., 2016; M. Singh et al., 2020). However, this compound was not observed as a potent molecule in *in-vitro* mycobacterium growth inhibition assay, which further warrants studies to improve potency. In general solubility and uptake of the drug molecules are the major factors for reaching therapeutic concentrations and achieving desired biological outcome.

To enhance the potency of this inhibitor we conjugated with a molecule which has been reported as cell penetrating in mammalian and bacterial cells. TAT peptides has been reported as cell penetrating peptide known to be used for enhancement of cellular entry and targeting of the drugs and bio-molecules (Deshayes et al., 2005). Previously, N-terminal tagging of peptides with different aromatic and aliphatic acids was employed for the enhancement of the potency of several molecules which have shown moderately antibacterial activity against the targeted pathogens (Bisht et al., 2007; Joshi et al., 2015; Laverty et al., 2010).

Apart from the cell penetrating efficiency of the TAT peptide it was also reported to exhibit the antimicrobial activity against bacteria, fungi and *Mtb* strains (Bhosle et al., 2018; Long Zhu & Shin, 2009; Wang et al., 2010). Although the antimicrobial mode of action of the cell penetrating peptides have been reported as multimodal and non-specific approaches due to their cationic nature, electrostatic interaction is possible with the anionic lipid membrane of bacteria (mimics of antimicrobial peptides) and intracellular targets due to their cell penetrating behaviour (Long Zhu & Shin, 2009; Rodriguez Plaza et al., 2014).

Targeted peptide conjugations or functionalized nano-formulations are usually efficient approaches to deliver the molecules to the target site. For example, cell-penetrating peptides have been extensively used to enhance the solubility and cellular uptake. TAT peptides are the most common moieties used to deliver chemotherapeutic agents which enhances cell permeability and also effective in killing off intracellular bacterial infections. It has been also shown that at concentrations used for targeted cellular delivery these peptides don’t exhibit toxicity towards mammalian cells.

In a similar approach we have conjugated a novel inhibitor (IN) with TAT peptide via cleavable (PC-1) and non-cleavable linkers (PC-2). For the cleavable linker the IN was conjugated via disulfide bond, which is labile inside mycobacterial cells due to mycothione reductase enzyme that is present in the cytoplasm (Patel & Blanchard, 1999). Therefore, the cleaved drug was expected to reach the DNA replisome at higher concentrations. PC-2 (conjugated with non-immolative linker) was directly synthesized on solid phase by conjugating IN with the N-terminal of the TAT peptide via amide bond. Due to the conjugation of IN with peptide the solubility was profoundly enhanced. Furthermore, the antitubercular efficacy of the conjugates was evaluated against *M. smegmatis* culture and found to be enhanced 2-8 fold compared to the non-conjugated molecule and TAT peptide alone.

The conjugate also exhibited inhibitory action against DnaG and Gyr in *in vitro* enzymatic assays at their MIC values. However, the TAT peptide was also able to inhibit the functions of the DnaG and Gyr enzymes at the observed MIC value of TAT in the identical conditions and this action probably originated from the electrostatic interaction of TAT peptide with DNA template/plasmid used in the assay. Previously, some cationic antimicrobial peptides (e.g. Buforin-II and Indolicidin) also have reported as inhibitors of genomic DNA or plasmids of the bacteria by electrostatic interaction and this has been proposed as an alternative mode of action of these peptides (Park et al., 1998; Subbalakshmi & Sitaram, 1998). However, to obtain more insight on this observation, further studies are warranted on the interactions of IN, its conjugates and TAT with enzymes by physical binding studies.

Biofilm development is an important factor for antimicrobial resistance, as it affords protection against antibiotics that are normally active against the same bacteria in its planktonic state. This antibiotic resistance of biofilm-forming microorganisms may result in treatment failure, and biofilms have to be physically eradicated to resolve the infection (Esteban & García-Coca, 2018). The effect of our experimental molecules, at their MIC ranges, on the formation of *M. smegmatis* biofilm was evaluated and significant inhibitory activity was noted.

However, the exact mechanism of the biofilm inhibition was not completely elucidated, although several studies reported that inhibition of DNA replication may cause the inhibition of biofilm formation (Trivedi et al., 2016). The unprecedented observation of the activity of the conjugates against biofilm formation inhibition of the *Msmg* gives an example of “moonlighting” peptides as reported previously (Rodríguez Plaza et al., 2012). Moonlighting peptides exhibit more than one activity in a single domain and differs from peptides that have multiple activities in different parts of peptide sequences (Rodríguez Plaza et al., 2012). Hence, IN and TAT peptide are moonlighting each other in conjugated form, but individually they were not efficient to inhibit the biofilm formation. A further study on cell cycle analysis of Msmg in biofilm culture is warranted to explore this observation.

Overall, we have demonstrated that inhibitor of bacterial replication, such as DnaG primase and gyrase inhibitor, as well as their TAT peptide conjugates, can efficiently inhibit growth of mycobacteria as well as biofilm formation. Therefore, this synthetic approach gives us a powerful tool to develop new and improved drug conjugates against *Mtb* in both planktonic and sessile forms.

## 4. Materials and Methods

### 4.1 Materials

All chemicals and solvents used for the synthesis and characterization of compounds, purchased from Sigma-Aldrich and Acros, were of reagent grade and used without further purification or drying. Chemicals used for biochemical assays were molecular-biology grade, purchased from Sigma-Aldrich. Ribonucleotides (ATP, CTP, GTP and UTP) were purchased from New England Biolabs (NEB). [α–32P] ATP (3000 Ci/mmol) and [γ–32P] ATP (3000 Ci/mmol) were purchased from Perkin Elmer. 1H NMR data were recorded on a 400 MHz Bruker NMR instrument. Chemical shifts were reported in δ (ppm) downfield from tetramethylsilane (TMS) and were recorded in appropriate NMR solvents (CDCl3, δ 7.26; CD3OD, δ 3.31; (CD_3_)_2_CO, δ 2.05; or other solvents as mentioned). All coupling constants (J) are given in Hertz. All NMR spectra were processed by MestReNova software. The following abbreviations are used: s, singlet; d, doublet; t, triplet; m, multiplet; br s, broad singlet peak. Mass spectra were recorded only for the molecules that exhibited best activity, using ESI-MS instrument. Thin layer chromatography (TLC) was performed on Merck aluminium backed plates, pre-coated with silica (0.2mm, 60F254) and the detection of molecules was performed by UV fluorescence (254 nm). Column chromatography separations were performed using silica gel (0.04-0.06 mesh) as stationary phase.

All the Fmoc protected amino acids and N,N,N’,N’-Tetramethyl-O-(1H-benzotriazol-1-yl)uronium hexafluorophosphate (HBTU) were purchased from Novabiochem and G.L. Biochem (Sanghai) Ltd. N,N,-Dimethylformamide (DMF) and dichloromethane (DCM) were purchased as biotech grade from Avantor J.T. Baker. Analytical HPLC was performed on a Dionex 1100 using a reverse phase C18 column at a flow rate of 1.5 mL/min. Preparative HPLC was performed on a Dionex Ultimate 3000 instrument using a C18 reverse phase preparative column at a flow rate of 20 mL/min. Mass spectrometry analysis was performed by LC-MS Thermo Surveyor 355.

### 4.2 Preparation of cell penetrating conjugates

Two strategies were employed to conjugate the DNA primase/gyrase inhibitor IN to the TAT peptide: i) conjugation of inhibitor via amide bond at the N-terminal of TAT sequence, on solid-phase resin (PC-2) and, ii) conjugation of inhibitor IN with TAT via mercaptoethanol linker (PC-1) (Figure 1). The PC-1 was prepared according to the synthesis scheme **2** (Figure 2). Briefly, the thiol group of 2-mercaptoethanol was protected with 2,2’-Dithiobis(5-nitropyridine) to form 2-((5-nitropyridin-2-yl)disulfanyl)ethanol (1a), which was further conjugated with IN via ester linkage to prepare the compound 1b. Furthermore, 1b was conjugated to the cysteine linked TAT peptide (NH_2_-CysTAT-CONH_2_) by solution phase synthesis through sulphide exchange reaction at pH-7.4 in 3-(N-morpholino)propanesulfonic acid (MOPS) buffer at room temperature. The conjugation reaction was monitored by RP-HPLC and ESI-MS (Figure S1-S8). The Fmoc based solid phase peptide synthesis strategy was used for the synthesis of TAT peptide (GRKKRRQRRRPQ-CONH2) on Rink amide MBHA resin by using an automated peptide synthesizer. All the coupling reactions were performed using HBTU coupling with 4 equivalents of the amino acid. For deprotection of Fmoc group, 25% v/v piperidine in DMF was used. Upon completion of peptide synthesis on resin, compound IN was conjugated to the N-terminal of last amino acid using HBTU and DIPEA for 4h. To cleave the resin bound conjugated peptides, a cleavage mixture comprising of trifluoroacetic acid/thioanisole/phenol/ethanedithiol/H_2_O (91.25:2.5:2.5:1.25:2.5) was used. Purification of the conjugated peptide was done on C-18 RP-HPLC column with linear gradients of water containing 0.1% TFA (gradient A) and acetonitrile. The final product was characterized by LC-ESI-MS (Thermo Surveyor 355).

**Figure: 2.**
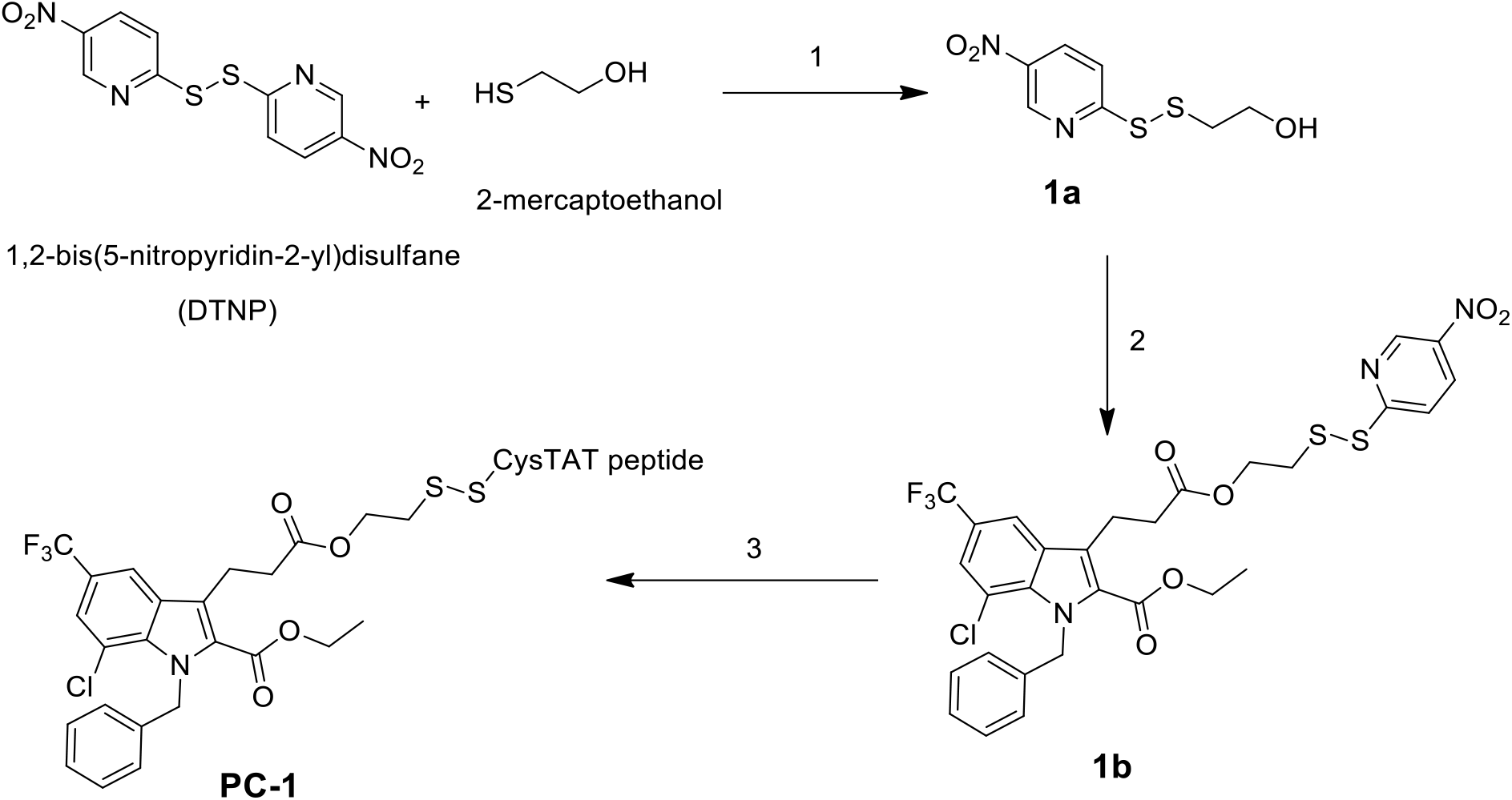
Synthesis scheme of sulphide linked peptide conjugates. Reagents and conditions: 1. DCM: methanol (50:50), 2hr 2. IN, DCC, DMAP in DMF, 18hr, 3. HS-CysTAT-peptide, MOPS buffer (pH 7.0): ACN (65%).

### 4.3 Anti-mycobacterium assay

*M*s*mg* mc^2^155 cells were grown in Middlebrook 7H9 broth (Fluka) liquid medium supplemented with glycerol (0.4%, v/v) and Tween-80 (0.05%, v/v) (7H9++ liquid medium). Solid medium was composed of Middlebrook 7H10 agar (Difco) and 0.4% (v/v) glycerol (7H10+ agar solid medium).

Anti-mycobacterium activity was evaluated using the broth micro-dilution method as reported previously (Khara et al., 2014). The inoculums were prepared from mid-log phase bacterial cultures. Each well of a 96-well polystyrene micro-titre plate was added 10 μL of serially diluted test compounds in 7H9 medium over the desired concentration range. Then 100 μL of ~10^5^ CFU/mL of bacterial suspension in 7H9 broth was added into each well of the plate except for the negative control well. The micro titre plates were incubated at 30 °C. After 48 h absorbance was read at 600 nm. Cultures (~10^5^ CFU/mL) without compounds were used as control. Un-inoculated broth was used as a negative control. Tests were carried out in triplicate. Minimum inhibitory concentration (MIC) is defined as the lowest concentration of test sequences that completely inhibited growth. For comparison standard antitubercular agent isoniazid (INH) was also assayed under identical conditions.

### 4.4 Oligoribonucleotide synthesis assay by *Mtb* DnaG primase

Standard 10 μL reaction contained 50 μM of DNA template (5’-CCGACCCGTCCGTAATACAGAGGTAATTGTCACGGT-3’), 250 μM GTP, CTP, UTP and [α-^32^P]-ATP, and *Mtb* DnaG primase in a buffer containing 40 mM Tris-HCl (pH 7.5), 1 mM MnCl2, 10 mM DTT, and 50 mM KGlu. After incubation at 37°C for 60 minutes, the reaction was terminated by adding an equal volume of sequencing buffer (98% formamide, 0.1% bromophenol blue, 20 mM EDTA). The reaction products were separated by gel-electrophoresis on a 25% polyacrylamide sequencing gel containing 7 M urea, and were visualized by autoradiography.

### 4.5 DNA supercoiling assays for *Mtb* DNA Gyrase

DNA supercoiling activity was tested with purified *M. tuberculosis* GyrA and GyrB subunits mixed in 2:1 ratio. The reaction mixture contained 50 mM HEPES [pH 7.9], 100 mM potassium glutamate, 6 mM magnesium acetate, 2 mM spermidine, 4 mM dithiothreitol, bovine serum albumin [50 μg/ml], 1 mM ATP and 13.34 ng/μl relaxed pBR322 DNA, 10% V/V DMSO. Gyrase proteins were added, and the reaction mixtures were incubated at 37°C for 1 h. Reactions were terminated by adding one volume of Stopping Buffer (40% sucrose, 100 mM Tris-HCl pH 8, 10 mM EDTA, 0.5 mg/ml Bromophenol Blue) and one volume of chloroform/isoamyl alcohol (v:v, 24:1). The reactions were vortexed, centrifuged briefly and loaded onto 1% agarose gel in 1X Tris-borate-EDTA buffer, pH 8.3 (TBE). Electrophoresis was performed for 4h at 40 V, and the gel was subsequently stained with ethidium bromide (0.5 mg/ml). One unit of enzyme activity was defined as the amount of DNA gyrase that converted 400 ng of relaxed pBR322 to the supercoiled form in 1 h at 37°C in 30 μl reaction.

### 4.6 Inhibition of biofilm formation of *Msmg*

Briefly, freshly grown *Msmg* mc^2^155 was diluted in fresh biofilm growth medium (Sauton’s medium) 100 times (100 μl culture was diluted in 10ml of medium). 1000 μl of diluted culture was dispensed in wells of a 24-well polystyrene plate for biofilm formation.

[Sauton’s media supplemented with 2% glucose preparations: Dissolve 0.5g of KH_2_PO_4_, 0.5 g of MgSO_4_, 4g of asparagine, 2g of Citric acid, 0,05g of ferric ammonium citrate, 60mL of glycerol in 900ml of water and add 2% w/v of glucose. Adjust the pH to 7.0 with NaOH. Autoclave, cool and just prior to starting the experiment, add sterile ZnSO_4_ to a final concentration of 0.1% w/v.]

### 4.7 Biofilm formation inhibition assay

To evaluate the inhibition of biofilm formation, compounds and Isoniazid at MICs and sub MICs were added initially with diluted culture. The plate was sealed with parafilm and incubated at 37 °C in an incubator for 7 days without shaking. After the 7th day of incubation 0.1 % Tween 80 was added to each well and gently swirled to dissolve the detergent. After 15 minutes of incubation pellicles were collected and washed thrice with PBS containing 10% glycerol and 0.05% Tween 80. Pellicles were kept on a rocker at 4 °C for 12 h in the same buffer for dispersal of matrix. Suspension was then homogenized by vortexing and diluted 10 times in buffer and 90μl of each sample was dispensed into 96 well polystyrene plates and 10 μl of alamar blue reagent was added to each well to check the viability of bacterial cells. The plate was further incubated at 37 °C for 3h and percentage reduction of alamar blue was calculated by measuring fluorescence by using a spectrofluorometer at excitation wavelength of 570nm and emission wavelength of 610nm. To further corroborate and validate the result of alamar blue assay the bacterial suspensions was diluted 10 times in PBS and 10 μl was plated on a MH10 agar plate for counting of colony forming unit (CFU).

## 5. Aknowledgements

This work was supported by grants from the Bi-National Science Foundation (BSF, grant number 2016142), the ISRAEL SCIENCE FOUNDATION (grant No. 1023/18), and the IMTI (TAMAT) / Israel Ministry of Industry-KAMIN Program, Grant No. 59081. We thank Prof. Gonen Ashkenazy for help with peptide synthesis. We are very thankful to Jack S. Cohen (Visiting Professor, BGU) for helpful discussions and language corrections in the manuscript.

## 6. Author contributions

R. P. D. and B. A. conceived and designed the study, R. P. D., M. S., S. I. and B. T. performed the experiments and analyzed the data, R. P. D. and B. A. wrote the manuscript with inputs from all the co-authors.

## 8. Data availability statement

The data that supports the findings of this study are available in the supplementary material of this article

