## Supplementary material for "Cell-penetrating peptide conjugates of indole-3-acetic acid-based DNA primase/Gyrase inhibitors as potent antitubercular agents against planktonic and biofilm culture of *Mycobacterium smegmatis*": Suplemental Figures and Tables

### ELECTRONIC SUPPLEMENTARY INFORMATION

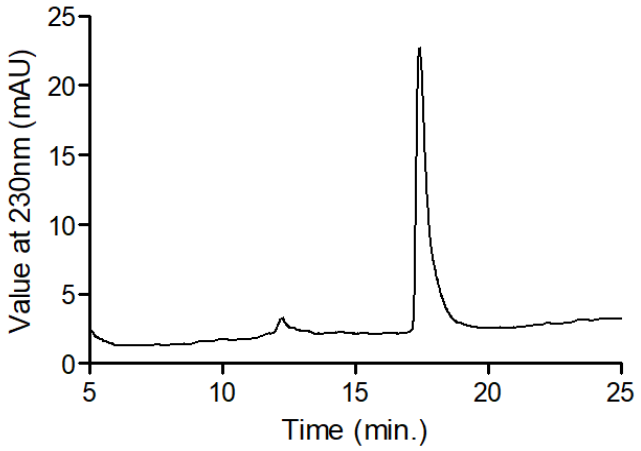


**Figure**: S1 HPLC profile of **PC-1**

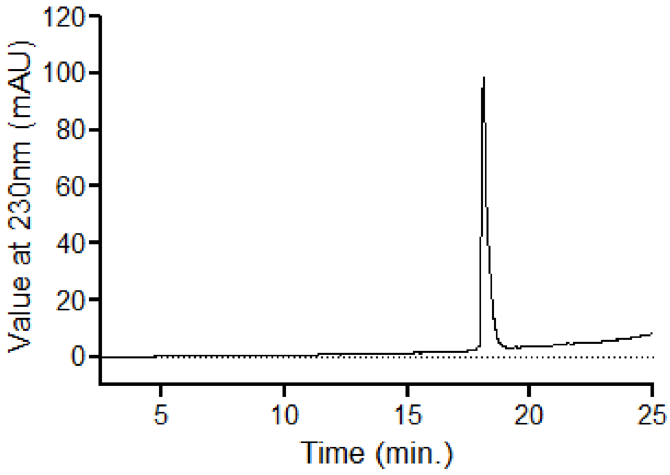


**Figure**: S2 HPLC profile of **PC-2**

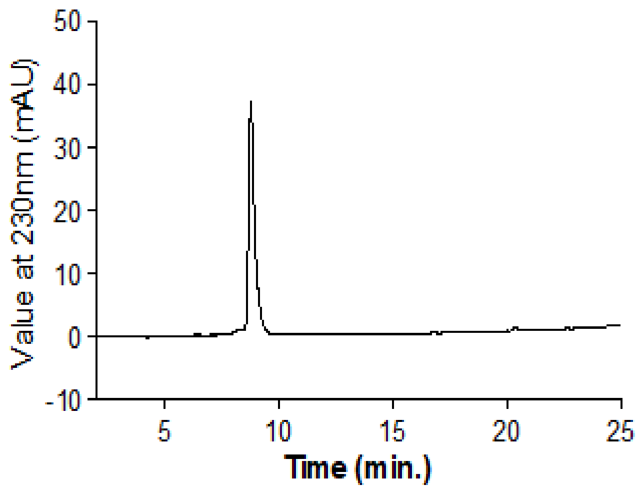


**Figure**: S3 HPLC profile of TAT peptide

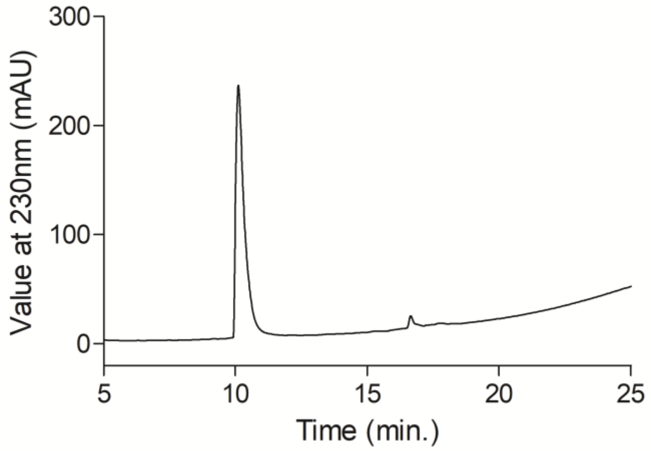


**Figure**: S4 HPLC profile of cysTAT peptide

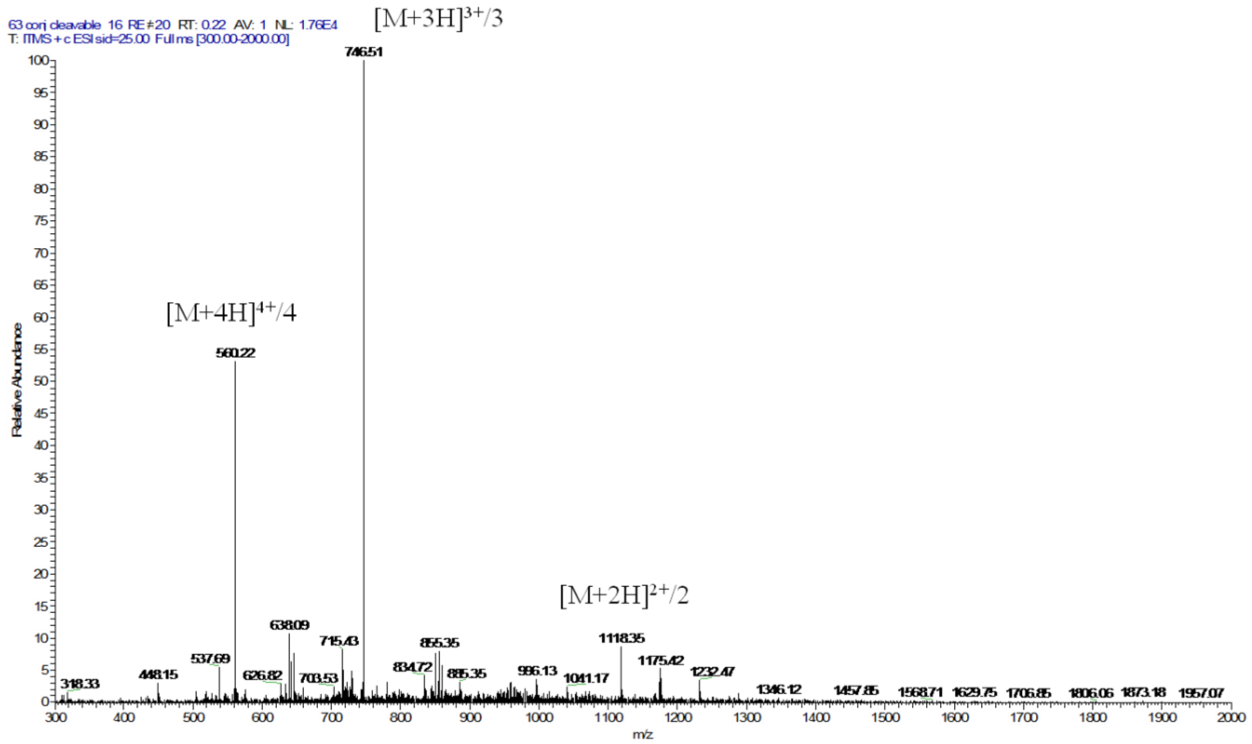


**Figure**: S5 ESI-MS spectrum of **PC-1**

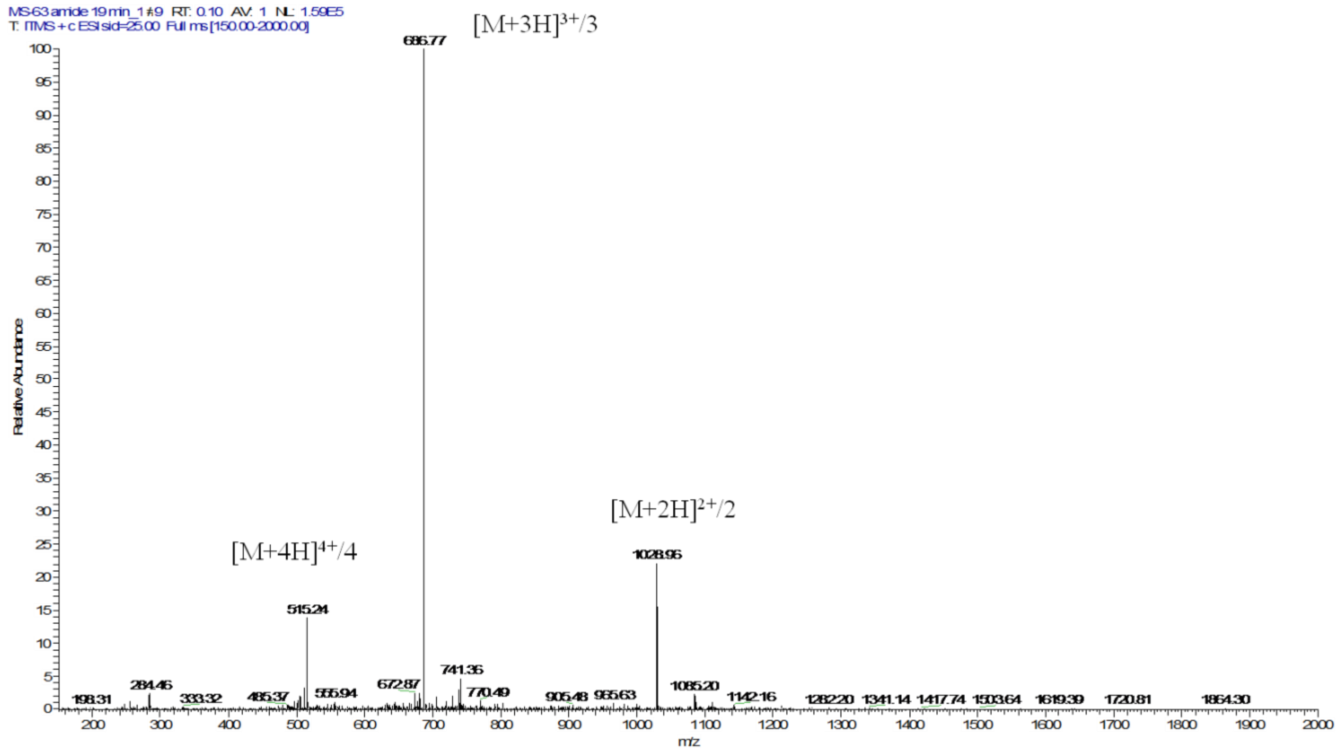


**Figure:** S6 ESI-MS spectrum of **PC-2**

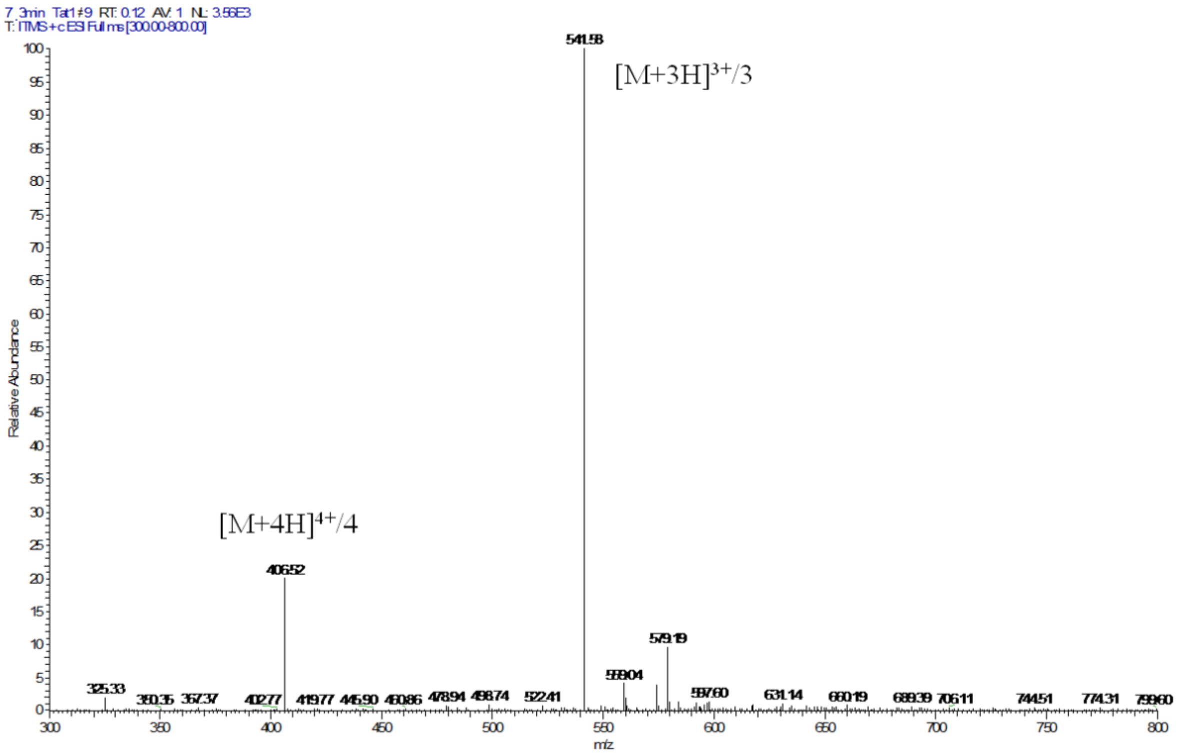


**Figure**: S7 ESI-MS spectrum TAT peptide

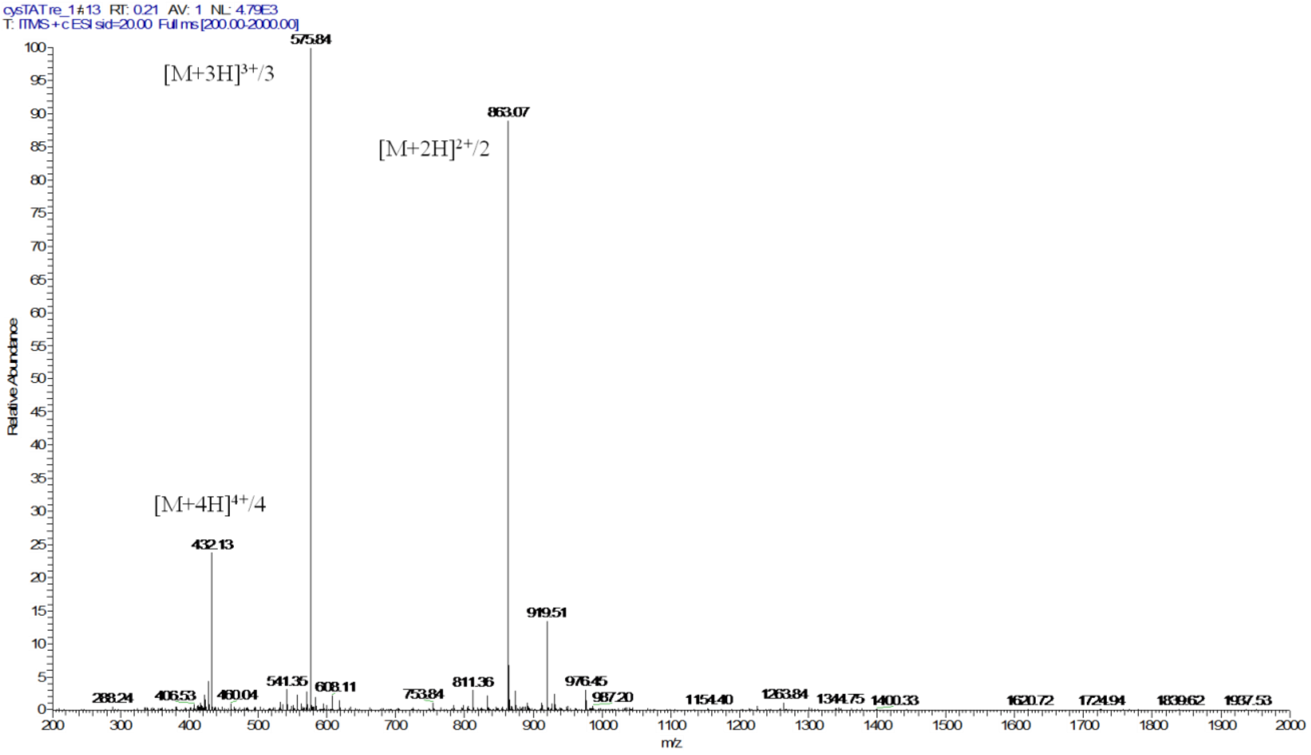


**Figure:** S8 ESI-MS spectrum cysTAT peptide

**Characterizations of Inhibitor and intermediate of linker:**

**3-(1-benzyl-7-chloro-2-(ethoxycarbonyl)-5-(trifluoromethyl)- 1H-indol-3-yl)propanoic acid** (**IN**) TLC (EtAc/Hex-40:60) Rf = 0.8; 25% yield; white solid; 1H NMR (400 MHz, MeOD) δ 8.09 (d, 1H), 7.53 (d, 1H), 7.27 – 7.13 (m, 3H), 6.82 (d, 2H), 6.26 (s, 2H), 4.33 (q, 2H), 3.39 (t, 2H), 2.63 (t, 2H), 1.31 (t, 3H); 13C NMR (101 MHz, MeOD) δ 176.5, 162.7, 140.8, 130.6, 130.5, 129.5, 128.0, 126.5, 126.2, 124.6, 124.5, 124.3, 119.3, 118.7, 118.7, 62.6, 50.1, 36.2, 21.7, 14.3; HRMS (ESI) calcd for $C_{22}H_{19}ClF_3NO_4$ for [M+ H] 454.1028, found 454.1027.

**2-((5-nitropyridin-2-yl)disulfanyl)ethanol** (**1a**), TLC (EtAc/Hex- 50:50), Rf = 0.6, 80% yield; white solid; $^1$H NMR (400 MHz, $CDCl_3$) δ 9.32 (d, *J* = 3.3 Hz, 1H), 8.36 (dd, *J* = 8.8, 2.6 Hz, 1H), 7.67 (d, *J* = 8.8 Hz, 1H), 4.09 (t, *J* = 6.8 Hz, 1H), 3.80 – 3.75 (m, 2H), 3.00 (t, *J* = 5.2 Hz, 2H); ESI-MS: m/z [M + H] calcd. for $C_7H_9N_2O_3S_2$: 233.00 Found: 233.06.

**Ethyl 1-benzyl-7-chloro-3-(3-(2-((5-nitropyridin-2-yl)disulfanyl)ethoxy)-3-oxopropyl)-5-(trifluoromethyl)-1H-indole-2-carboxylate** (**1b**)

TLC (Methanol/DCM 20:80) Rf = 0.78; 92% yield; white solid; $^1$H NMR (400 MHz, $CDCl_3$) δ 9.29 (dd, *J* = 2.6, 0.7 Hz, 1H), 8.41 (dd, *J* = 8.8, 2.6 Hz, 1H), 7.94 (dd, *J* = 1.6, 0.8 Hz, 1H), 7.84 (dd, *J* = 8.9, 0.7 Hz, 1H), 7.53 (dd, *J* = 1.7, 0.6 Hz, 1H), 7.28 – 7.21 (m, 3H), 6.90- 6.89 (m, 2H), 6.28 (s, 2H), 4.36 (q, *J* = 7.1 Hz, 2H), 4.33 (t, *J* = 6.5 Hz, 2H), 4.17 (d, *J* = 7.4 Hz, 2H), 3.41 (t, *J* = 7.4 Hz, 2H), 3.02 (t, *J* = 6.4 Hz, 2H), 2.71 (t, *J* = 7.2 Hz, 2H), 1.33 (t, *J* = 7.2 Hz, 3H); ESI-MS calcd for $C_{29}H_{25}ClF_3N_3O_6S_2$ for [M+ H] 668.08, found 668.09.

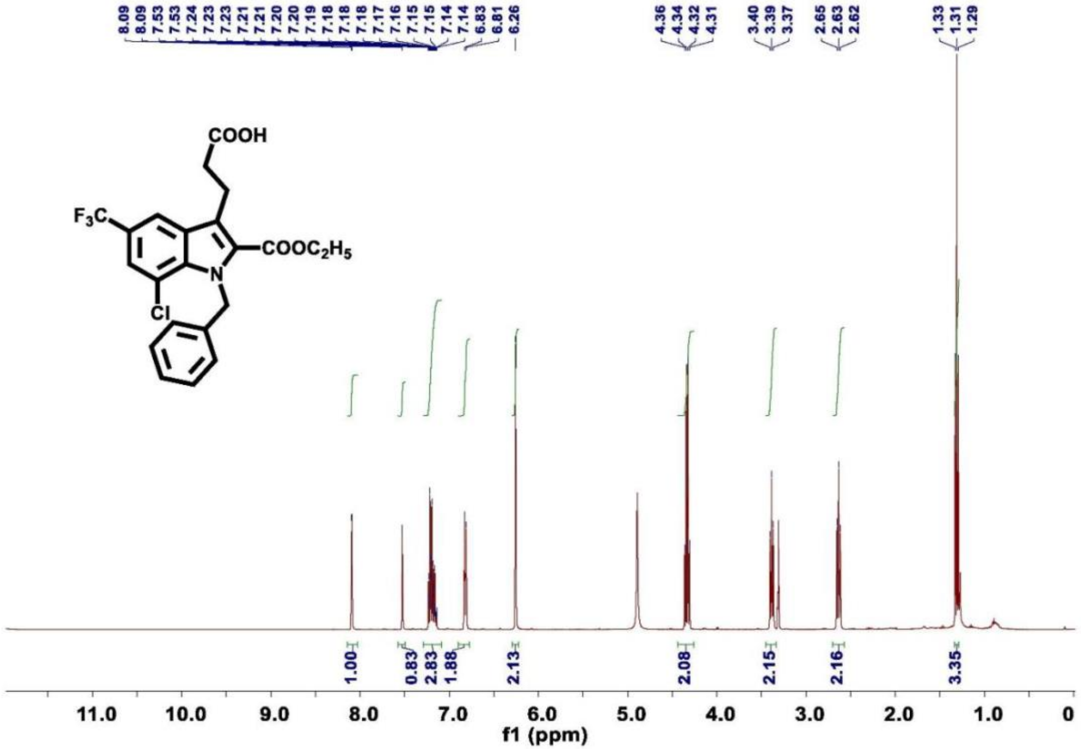


**Figure**: S9 $^1$H NMR of compound IN

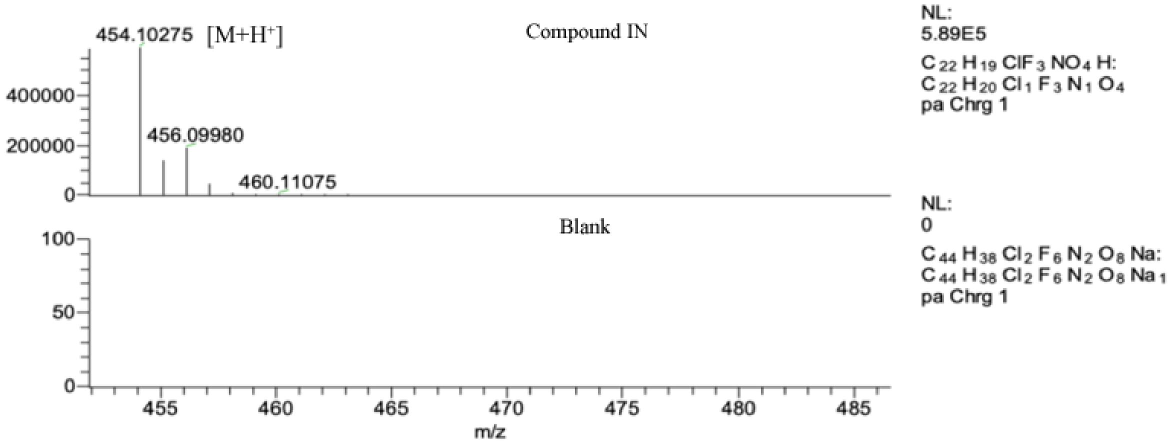


Figure S10: HR-ESI-MS of compound IN

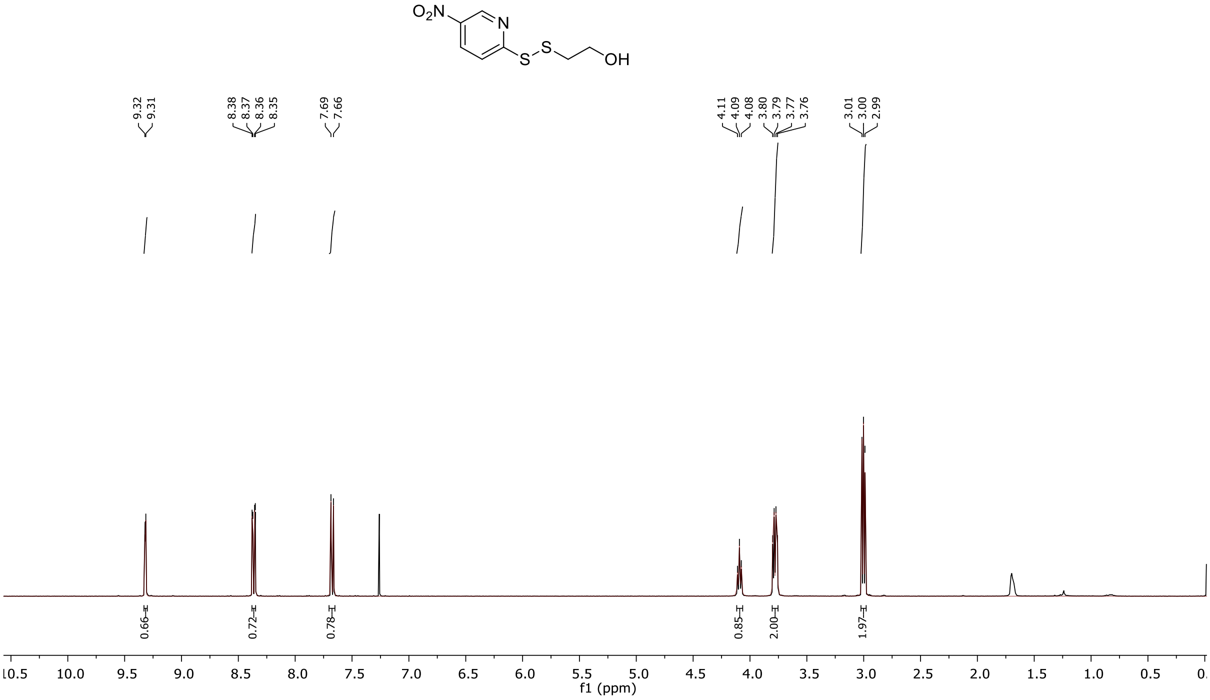


**Figure**: S11 $^{1}$H NMR of compound 1a

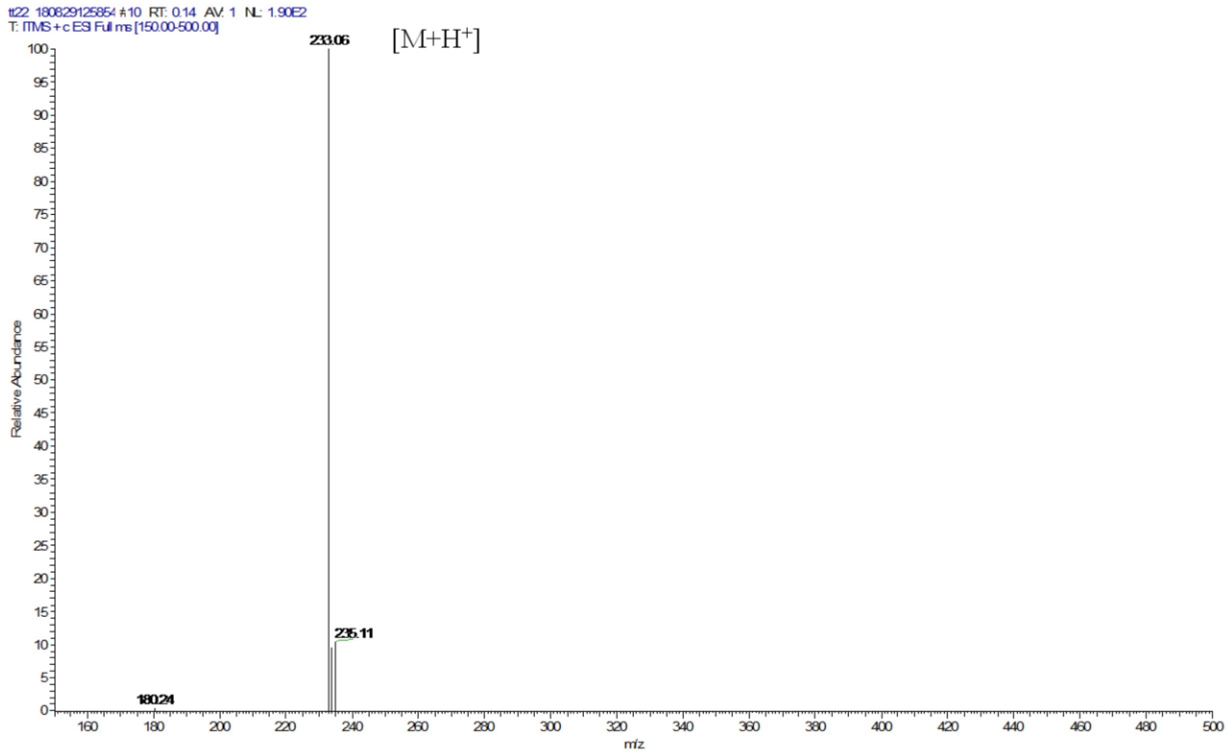


**Figure**: S12 ESI-MS of compound 1a

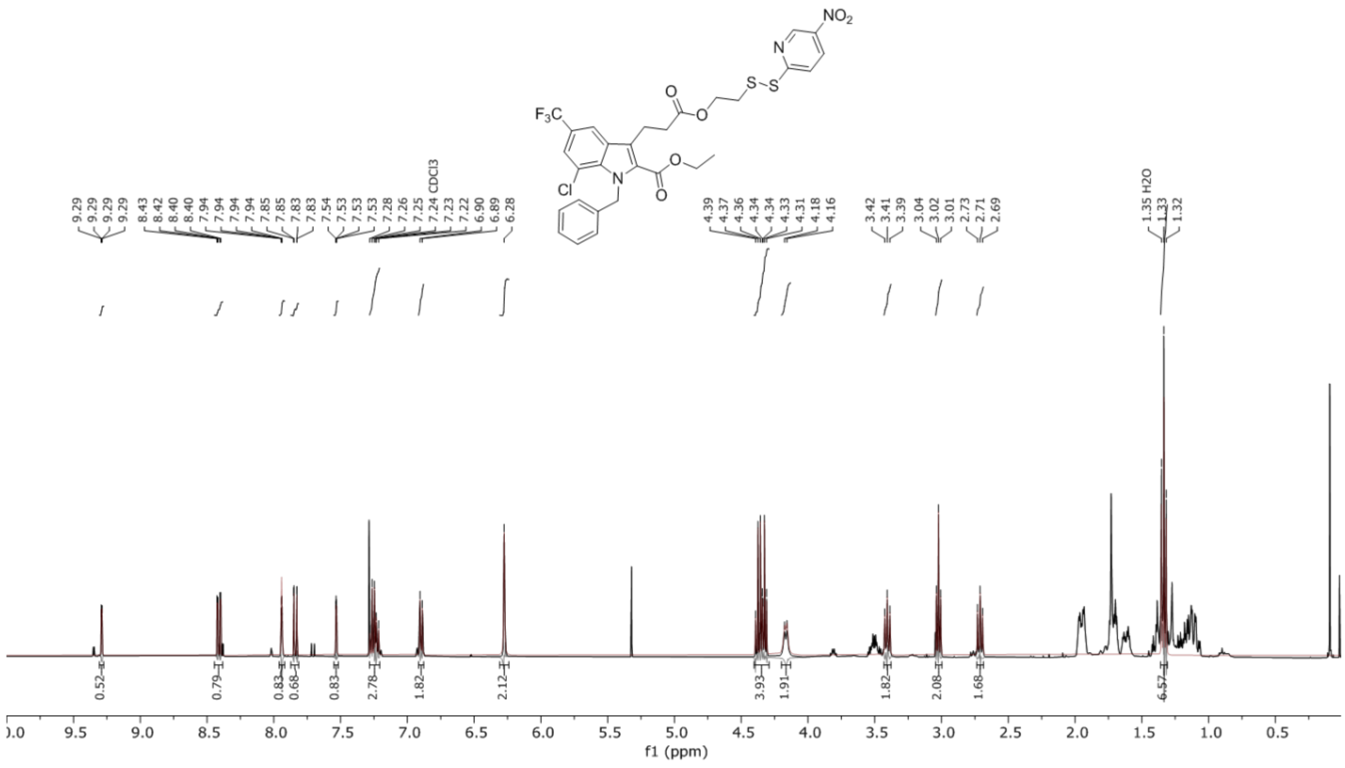


**Figure**: S13 $^{1}$H NMR of compound 1b

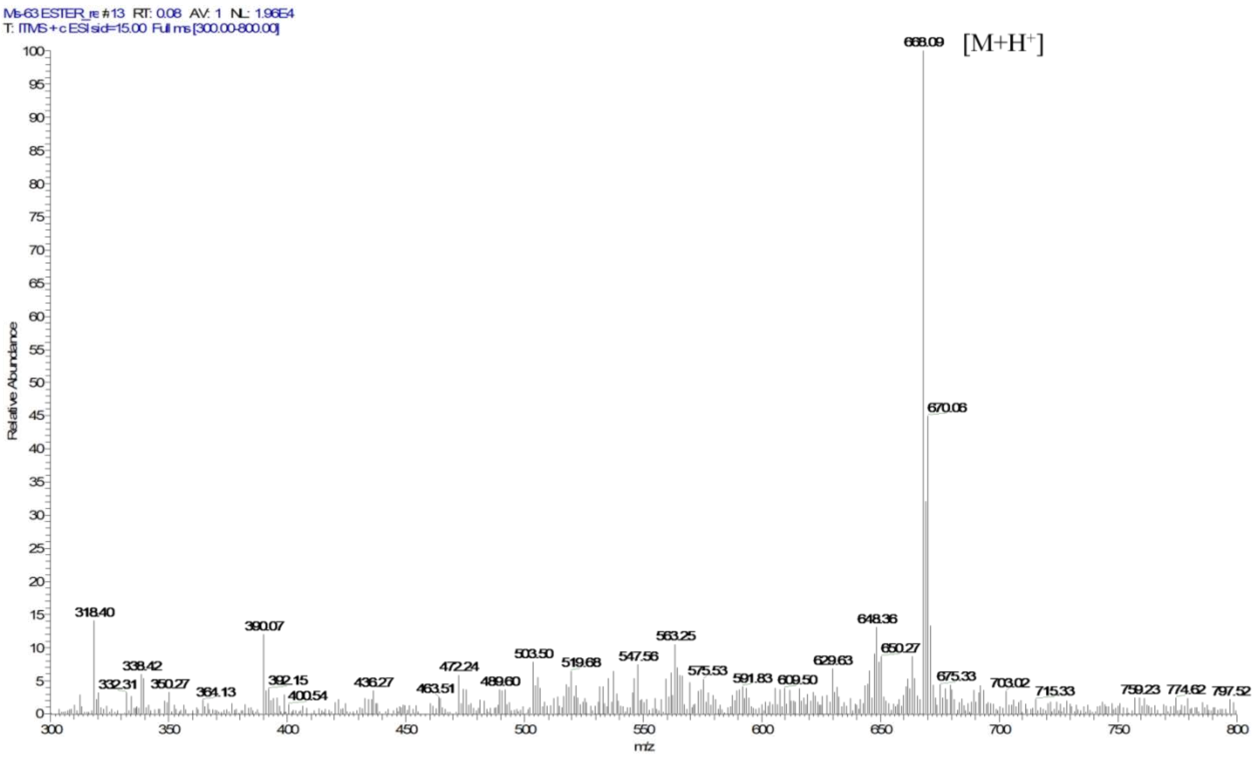


**Figure:** S14 ESI-MS of compound 1b

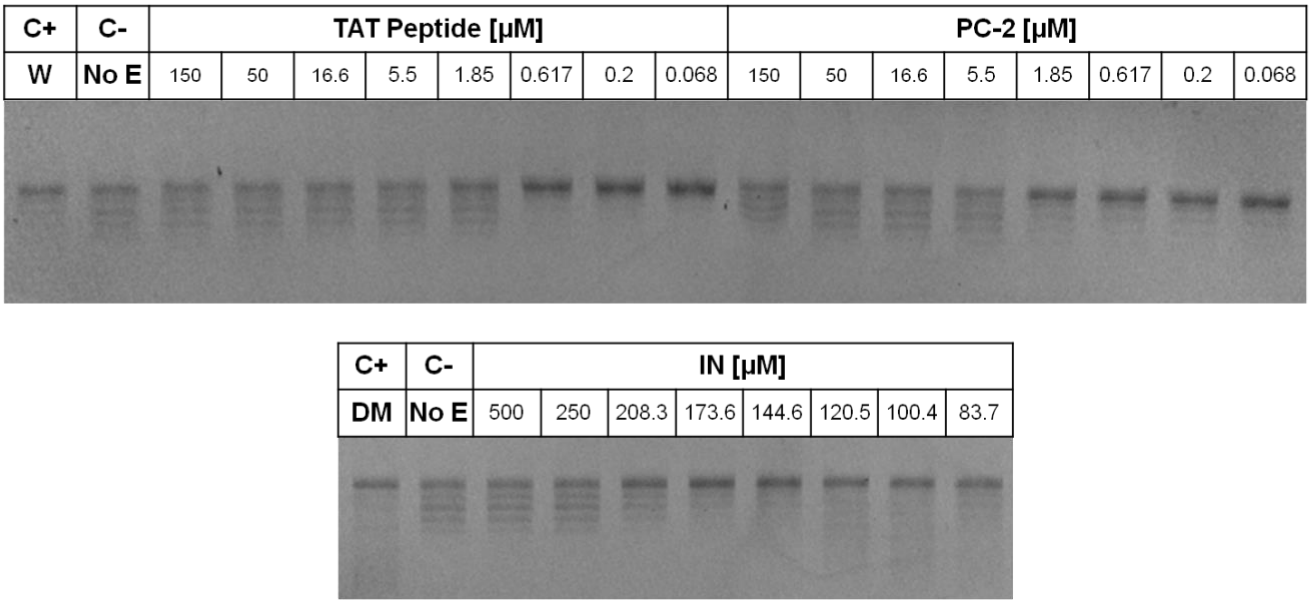


**Figure**: S15 Effect of inhibitor molecule (IN), PC-2 and TAT peptide on inhibition of DNA supercoiling by small molecule inhibitors. The 20µL reaction containing Mtb-DNA Gyr (19.5 nM), 13.24ng/µL pBR322 DNA and 1mM ATP in the presence of inhibitors was incubated at 37 °C for 60 min. The reaction products were resolved in 1% agarose gel followed by staining with EtBr.
